# The contribution of the European LIFE program to mitigate damages caused by large carnivores in Europe

**DOI:** 10.1101/2021.06.25.449866

**Authors:** Teresa Oliveira, Adrian Treves, José Vicente López-Bao, Miha Krofel

## Abstract

Governments around the world invest considerable resources to reduce damages caused by large carnivores on human property. To use these investments more efficiently and effectively, we need to understand which interventions successfully prevent such damages and which do not. In the European Union, the LIFE program represents by far the largest financial instrument to help EU Member States with the implementation of conservation activities, including mitigation of damages caused by large carnivores. However, we currently lack information about the effectiveness of this funding program in reducing carnivore damages. We reviewed 135 LIFE projects dealing with large carnivores between 1992 and 2019 to provide an overview of the use of damage prevention methods and evaluate their functional and perceived effectiveness. Methods evaluated ranged from non-lethal and lethal interventions, to information dissemination and compensation schemes. The largest number of the projects was focused on grey wolf (*Canis lupus*) and brown bears (*Ursus arctos*) in the Mediterranean countries and in Romania. Electric fences were reported as the most successful method for reducing damages by large carnivores, and most of the non-lethal methods used showed at least moderate effectiveness. However, standards of measuring and reporting effectiveness were in general relatively low, which limits our ability to measure actual impact. Therefore we urge project managers and evaluators to improve these standards, as well as the dissemination of the project results. We provide a list of recommendations for improving measuring and reporting success of implemented interventions for the benefit of future projects aimed to reduce damages caused by wildlife.

**Article impact statement:** Electric fences were reported as the most effective method to prevent large-carnivore damages and are recommended for future use.

## Introduction

The predatory behavior of large carnivores, threatening livestock, pets and, sometimes, human safety, often represent the main factor opposing the landscape-sharing approach to coexist with these species (López-Bao et al., 2017; Treves & Karanth, 2003; van Eeden et al., 2018; Woodroffe et al., 2005). In Europe, the last decades were marked by a recovery of large carnivore populations (Chapron et al., 2014) which in many areas lead into an increase of damage caused to livestock and other human properties (Bautista et al., 2019; Fernandez-Gil et al., 2016; Hovardas, 2018; Musiani et al., 2003). Areas where large carnivores have been absent for several decades or centuries are often particularly sensitive to conflict situations, due to lack of experiences in coexistence with predators and disuse of traditional methods to prevent damages (Salvatori & Mertens, 2012; López-Bao et al., 2017). Therefore, the recolonization of former territories is raising challenging conservation issues, including damages, fear for personal safety, as well as conflicts between different interest groups on how to conserve large carnivores (Boitani & Linnell, 2015; Hovardas & Marsden, 2018; Treves et al., 2004). While some groups are concerned with carnivore conservation and welcome their recolonization, others (e.g. farmers and livestock owners) are distressed due to damages they already suffered or may suffer (Hovardas, 2018) and the need for additional work required for preventing these damages. Nevertheless, despite several different interests, there is often a common goal towards which all of these groups strive, i.e. reduction of human-carnivore conflicts.

Governments around the world invest considerable logistical and budgetary resources in finding win-win solutions for carnivore restoration and long-term preservation at the same time as protecting agricultural incomes and safety of domestic animals and people (Carter & Linnell, 2016; Chapron et al., 2014; Ripple et al., 2014; Van Eeden et al. 2018). Those investments would be used more efficiently and effectively if we knew which interventions achieved which of those two goals, if any, and how they did so (Treves et al. 2019). Several damage mitigation methods (hereafter, DMMs) are applied with the attempt of reducing damages caused by large carnivores and facilitating the coexistence between carnivores and humans (Dickman, 2010; Woodroffe et al., 2005). These methods can be generally grouped in two main categories: reactive and proactive. Reactive DMMs include, for example, a large part of damage compensations schemes aimed to mitigate the economic burdens for livestock owners after they report an attack (e.g., Bautista et al., 2019; Giannuzzi-Savelli et al., 1997; Ravenelle & Nyhus, 2017), while proactive DMMs aim at preventing damages before they occur, such as electric fences and the use of carnivore-proof garbage bins, to reduce the access to anthropogenic food by carnivores (Linnell et al., 1996; Shivik et al., 2003; Van Eeden et al., 2018). However, due to the high variety of socio-ecological contexts in which damages occur, each situation should be considered carefully before applying any type of DMM (Hovardas & Marsden, 2018; Treves et al., 2009; van Eeden et al., 2018).

The choice of a DMM to be implemented should primarily rely on its effectiveness for achieving an explicit outcome (Treves et al., 2016, 2019; van Eeden et al., 2018). Effectiveness of DMM can be assessed from two stand-points: perceived and functional. While the first considers qualitative self-assessments (i.e., subjective perception of effectiveness of given practice (Ohrens et al., 2019; Treves et al., 2019), the latter considers a quantitative and measurable approach (i.e., an actual measured reduction or increment in damages, retaliations, or other physical manifestation of conflicts; Treves et al., 2016, Eklund et al. 2017). Since perceived effectiveness is less reliable and can differ considerably from functional effectiveness (Ohrens et al. 2019), relying solely on perceived effectiveness might lead to a suboptimal choice of DMM (Treves et al., 2009, 2019). However, despite the fact that numerous academic studies, conservation projects and management strategies recommend or implement DMMs, our knowledge on their functional effectiveness under different conditions is still surprisingly limited (Eklund et al., 2017; Miller et al., 2016; Treves et al., 2019, 2016; Van Eeden et al., 2018). A careful evaluation of the effectiveness of each DMM applied, as well as a clear and unbiased reporting of the results, is essential to obtain sound information on the best method to be implemented in each context, and to create particular guidelines. This is especially important for funding programs that aim to support coexistence between people and large carnivores at large spatial scales (Van Eeden et al. 2018).

In 1992, the European Commission initiated the LIFE program (*L’Instrument Financier pour L’Environnement;* https://ec.europa.eu/easme/en/life) as a financial instrument to help EU Member States with the implementation of conservation actions. It is currently by far the largest funding program dedicated to nature and environment (€5.45 billion is planned for the next 6-year cycle, 2021-2027; https://ec.europa.eu/easme/en/section/life/life-history-life). A large part of the program is dedicated to conserving and managing habitats and species, according to the EU’s directives on birds and habitats, and gives priority to strictly protected species (e.g., Annex IV in the Habitats Directive) and the Natura 2000 network (https://ec.europa.eu/environment/nature/index_en.htm).

The three largest and most widespread species of carnivores in Europe (grey wolf, *Canis lupus*; Eurasian lynx, *Lynx lynx*; brown bear, *Ursus arctos*) are listed as priority species for the LIFE program and feature prominently among the species aimed by the LIFE projects. Among them, many have implemented a broad number of interventions aimed at mitigating human-carnivore conflicts, both proactive and reactive, from the distribution of electric fences to outreach activities (see below). Although LIFE projects started with projects conducted at national level, there has been steady increase in transboundary projects (e.g., Salvatori & Mertens, 2012) because most large carnivore populations in Europe are transboundary (Boitani et al., 2015; Chapron et al., 2014; Linnell et al., 2008). So far, €88 million have already been provided to projects that included mitigation of damages caused by bears, wolves and lynx (plus additional €36 million granted to ongoing projects). However, although large funds have been invested, we currently lack information about the effectiveness of the interventions implemented under this funding program to reduce human-carnivore conflicts. Without such knowledge, it is difficult to evaluate the success of the LIFE program in improving human-carnivore coexistence, as well as to provide recommendations for future efforts.

Here, we reviewed the completed and on-going LIFE projects dealing with large carnivore species in Europe over the past 28 years (1992-2019) with the aim to i) synthesize the use of DMMs within LIFE projects, ii) evaluate the trends in the promotion of DMMs within this program (by carnivore species and biogeographical area) and iii) evaluate the perceived and functional effectiveness of the interventions adopted in reducing human-carnivore conflicts. To our knowledge, this is the first study to provide a critical evaluation of the contribution of the LIFE program to reduce human-carnivore conflicts across Europe. Based on our results and information available, we developed recommendations for future efforts to facilitate human-wildlife coexistence, and to improve the standards of measuring the functional effectiveness of these interventions. We also identify deficiencies in reporting on effectiveness of implemented DMM and provide recommendations for improving dissemination of results for the benefit of the future projects and research.

## Methods

### Review of Projects

We compiled a database with projects from the LIFE’s database containing all completed and on-going projects between 1 January 1992 and 30 June 2019 (http://ec.europa.eu/environment/life/project/Projects/index.cfm). The search included the subtheme “Mammals” within the theme “Species”. We manually checked the summary of each LIFE project and selected those that mentioned any species of large carnivores and noted whether the project included implementation of at least one DMM. When the project summary was ambiguous about the use of DMMs, we checked the final report or a similar document, as well as the project’s webpage (when existing), to verify whether DMMs were used or not. We first retrieved general information about the project: 1) leading country, 2) target carnivore species, and 3) whether DMMs were implemented.

Then, for the projects including DMMs, we extracted information about 4) the type of DMM considered (Appendix S1), 5) effort, 6) functional and 7) perceived effectiveness (see below for details). As effort we considered the investment in the DMM (e.g. in terms of time, equipment, damage objects, costs), which can be relevant for comparisons of effectiveness among projects. For this section, we considered only completed projects. Although we first included all large carnivore species in Europe in our search, in the end we only evaluated projects including brown bears, grey wolves, and Eurasian lynx, because we found no completed LIFE project involving wolverine (*Gulo gulo*), and the reports of the projects involving the Iberian lynx (*Lynx pardinus*) did not include any DMM (probably because this species is rarely associated with damages caused to human property; Garrote et al., 2013).

### Categorization of mitigation practices

We first randomly selected a subset of projects (*n* = 35) and screened the final project reports to the EU, to identify the most commonly used DMMs. Based on this initial screening, we defined the following categories of DMMs (definition for each category is provided in Appendix S1): Large carnivore emergency teams (LCET), Information dissemination, Damage compensation schemes, Visual and sound deterrents, Electric fences, Physical barriers, Preventing access to anthropogenic food sources, Improvement of agricultural/farming practices, Increasing food availability for carnivores, Livestock guarding dogs (LGDs), Livestock guarding people (Shepherds), and Predator removal. In some cases, when the final report was not available, the goal of the information dissemination activities was not clear (i.e. we could not determine whether it was limited to education about carnivore ecology, status, conservation etc., or if it also included information about conflict prevention and DMMs applied within the project). In these cases, if the project summary mentioned the use of information dissemination and DMMs were conducted within the project, we assumed that those activities included dissemination of information on DMMs. For damage compensation schemes, we only considered cases in which the project was somehow related to this action (e.g. the project paid the compensation or if it changed or adapted the compensation scheme in the project area).

### Evaluating functional and perceived effectiveness

The methods used for the evaluation of effectiveness per DMM varied considerably among projects, thus data was not collected in a standardized way. For example, some projects reported only damage reduction in terms of number of animals killed before and after the implementation of an intervention, while others only reported the reduction in number of sheep folds affected. This variability in methodology prevented us from being able to directly compare effectiveness of different DMMs or to explore the relationship between effectiveness and additional factors, such as effort invested. Therefore, we conducted a two-step categorization to measure functional and perceived effectiveness of each DMM.

First, we extracted the reported functional and perceived effectiveness for given DMM from each project report. For data on functional effectiveness, we only considered reports, where comparison with a time period before the implementation was provided, i.e. before-and-after comparison also known as a silver standard of evidence (Treves et al., 2019). Functional effectiveness was usually provided as a value (e.g., the attacks in a given pasture were reduced by 55% after the deployment of electric fences compared to same period before the project), whereas perceived effectiveness was mostly reported qualitatively by the beneficiaries (e.g., owners considered electric fences to be very effective at reducing attacks in treated pastures). To enable comparison, we categorized the quantitative data in four categories: non-effective (no detectable difference in before-and-after comparisons), 1–25% reduction in damages caused by large carnivores, 26–75% reduction, and 76–100% reduction. For perceived effectiveness, we used the following qualitative classes based on descriptions available: none, medium, high, and very high effectiveness.

Secondly, we assessed the overall perceived and functional effectiveness of each DMM used in LIFE projects (all projects pooled). If more than 50% of the projects using a given DMM had reported 76-100% functional effectiveness for a given DMM and none reported it to be non-effective or counterproductive, then we classified that method as “Highly Effective”. A similar approach was applied when evaluating perceived effectiveness, where a DMM was considered to be perceived as “Highly Effective” when it was evaluated as having very high effectiveness in more than 50% of the projects and in none as being not-effective. “Effective” methods were those evaluated as “76-100%” or “very high” in ≤ 50% of the projects, and the remaining as having less than 75% functional effectiveness or “medium” and “high” perceived effectiveness, respectively. When more than 50% of the projects found given DMM to be non-effective in terms of functional or perceived effectiveness, we classified it as “Not Effective”. We classified the method as having “mixed results”, if a single project had reported a given DMM as having non-effective functional effectiveness (or “none” perceived effectiveness), while the other projects reported it to have higher functional (or perceived) effectiveness.

## Results

### Overview of the use of damage mitigation methods within LIFE projects

We retrieved information from a total of 263 LIFE projects focused on mammals, and, among them, more than half involved at least one species of large carnivore (n = 135). Most LIFE projects targeting large carnivores included at least one DMM (64%, n = 87, Fig. 1). From these, 74 projects were concluded by June 2019, while the remaining 13 were still in progress.

**Figure 1.**
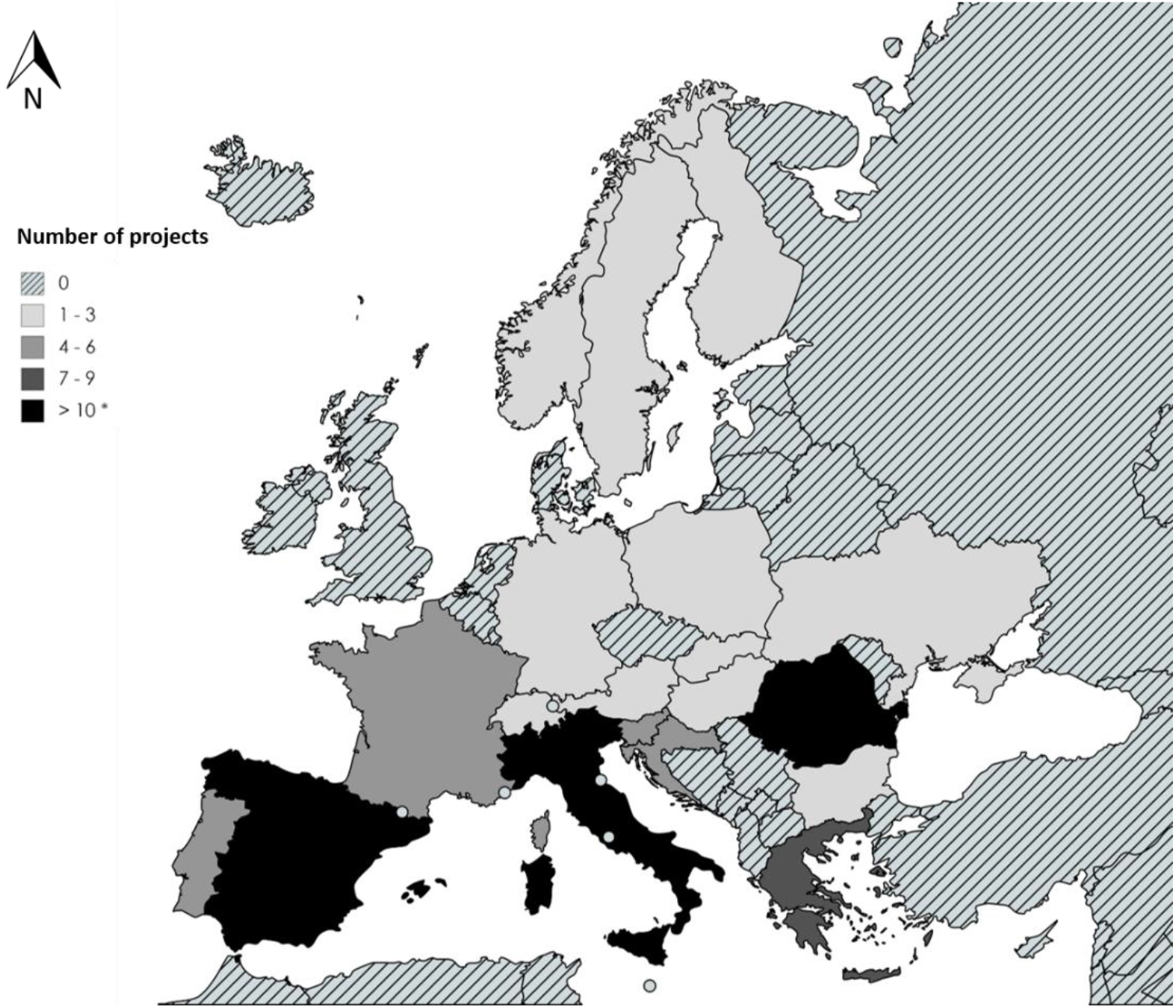
Number of LIFE projects including damage mitigation methods (DMM) per country, concluded and ongoing (n = 87). * The number of projects is much higher for Italy and Spain (n=25, for each of them)

Overall, the most frequently used DMM was dissemination of information to the stakeholders (conducted in 92% of the projects), followed by implementing electric fences (either fixed or mobile), which were used in 47% of the projects (Table 1). Other DMMs included, in decreasing order of use: improvement or implementation of damage compensations schemes, use of livestock guarding dogs (LGDs), and increasing food availability for carnivores, which were applied in 30-50% of the projects. The remaining DMMs were used infrequently, i.e. with < 25% of projects applying them (Table 1).

**Table 1.** Number of completed projects including given damage mitigation method (DMM), per species and overall, as well as the proportion of projects including given DMM (%).

| DMM | Grey Wolf |  | Brown Bear |  | Eurasian Lynx |  | Overall |  |
| --- | --- | --- | --- | --- | --- | --- | --- | --- |
|  | N<sub>proj</sub> = 34 |  | N<sub>proj</sub> = 59 |  | N<sub>proj</sub> = 5 |  | N<sub>proj</sub> = 74 |  |
|  | N<sub>proj</sub> | % | N<sub>proj</sub> | % | N<sub>proj</sub> | % | N<sub>proj</sub> | % |
| Information dissemination | 32 | 94.12 | 50 | 84.75 | 5 | 100.00 | 68 | 91.89 |
| Damage Compensation | 14 | 41.18 | 26 | 44.07 | 1 | 20.00 | 32 | 43.24 |
| Electric Fences | 14 | 41.18 | 28 | 47.46 | 4 | 80.00 | 35 | 47.30 |
| Improvement of agricultural practices | 1 | 2.94 | 6 | 10.17 | 1 | 20.00 | 6 | 8.11 |
| Increasing food availability | 7 | 20.59 | 21 | 35.59 | 1 | 20.00 | 25 | 33.78 |
| LCET | 0 | 0.00 | 10 | 16.95 | 0 | 0.00 | 10 | 13.51 |
| Livestock guarding dogs (LGD) | 12 | 35.29 | 13 | 22.03 | 1 | 20.00 | 22 | 29.73 |
| Physical barriers | 3 | 8.82 | 11 | 18.64 | 0 | 0.00 | 14 | 18.92 |
| Predator removal | 1 | 2.94 | 1 | 1.69 | 0 | 0.00 | 2 | 2.70 |
| Preventing access to anthropogenic food sources | 1 | 2.94 | 6 | 10.17 | 0 | 0.00 | 6 | 8.11 |
| Shepherds | 3 | 8.82 | 5 | 8.47 | 0 | 0.00 | 8 | 10.81 |
| Visual and sound deterrents | 1 | 2.94 | 1 | 1.69 | 0 | 0.00 | 2 | 2.70 |

**Table 2.**
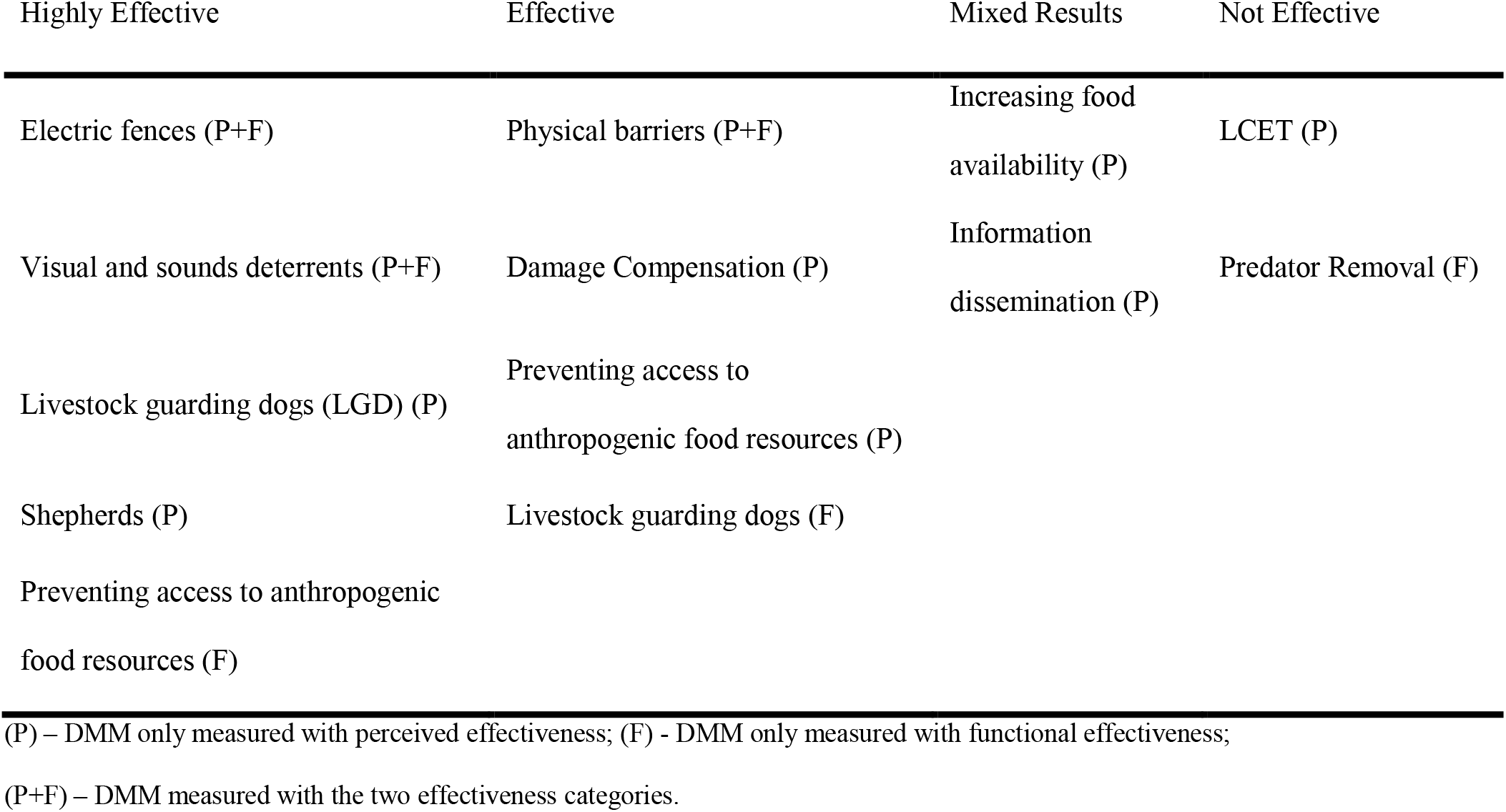
Evaluation of damage mitigation methods (DMMs) based on perceived and functional effectiveness.

| Highly Effective | Effective | Mixed Results | Not Effective |
| --- | --- | --- | --- |
| Electric fences (P+F) | Physical barriers (P+F) | Increasing food availability (P) | LCET (P) |
| Visual and sounds deterrents (P+F) | Damage Compensation (P) | Information dissemination (P) | Predator Removal (F) |
| Livestock guarding dogs (LGD) (P) | Preventing access to anthropogenic food resources (P) |  |  |
| Shepherds (P) | Livestock guarding dogs (F) |  |  |
| Preventing access to anthropogenic food resources (F) |  |  |  |

(P) – DMM only measured with perceived effectiveness; (F) - DMM only measured with functional effectiveness;
(P+F) – DMM measured with the two effectiveness categories.

We observed similar general patterns when individual species were considered separately. Notable exceptions were the development of LCETs and bear-proof garbage bins for preventing conflicts with bears (17% and 10% of the LIFE projects targeting bears, respectively; Table 1). As only few of the concluded projects included Eurasian lynx, and all of them included also other large carnivore species, it was not possible to see any pattern particular to lynx in the frequency of DMM use (Table 1).

#### Effectiveness of damage mitigation methods

We observed a surprising lack of information about the functional and perceived effectiveness of DMM, which were reported only for 14% and 31% of the concluded projects involving DMM (n= 74), respectively. Also, data on the effort spent for their implementation was limited (completely lacking in 39% of project reports), and differed significantly among projects (e.g. number of electric fences distributed per sheep fold or only number of sheep folds that got electric fences). Therefore, we could not compare effort and its influence on effectiveness in subsequent analyses.

#### Functional effectiveness

Information on functional effectiveness was not available in any of the reports for several of the DMMs (LCET, information dissemination, damage compensation schemes, improvement of agricultural practices, increasing food availability for carnivores, shepherds), and it was most often reported for electric fences (n = 13; Fig. 2).

**Figure 2.**
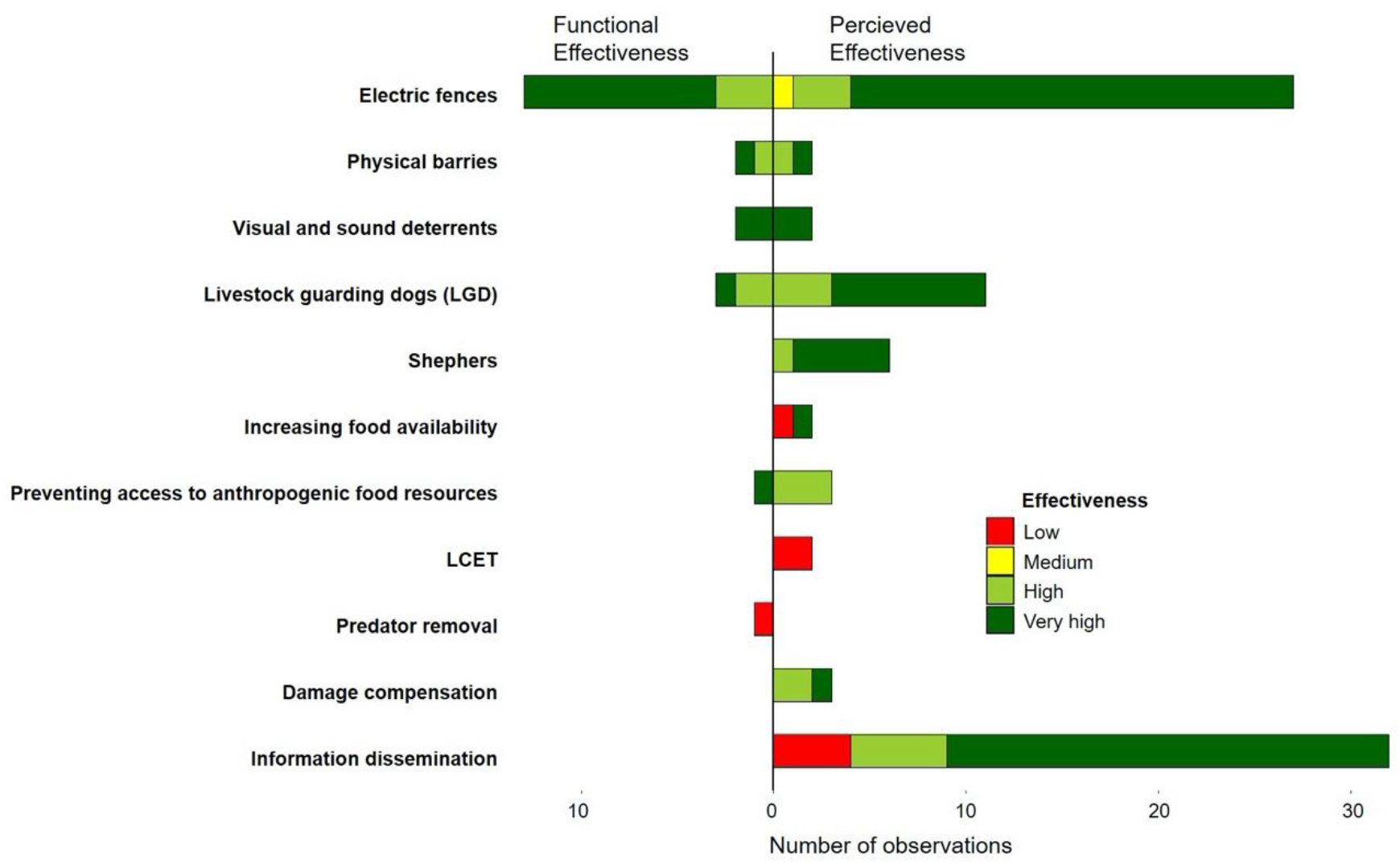
Measure of functional and perceived effectiveness (quantitative and qualitative assessments, respectively) of each damage mitigation method (DMM). An observation corresponds to a DMM measured within a project.

Electric fences appear to be the most effective DMM for reducing damages according to the available information; in most of the reported cases their effectiveness was >75% (i.e. highly successful; Fig. 2). High success was reported also for physical barriers, visual/sound deterrents, and preventing access to anthropogenic food resources, although sample sizes were much lower for these DMMs (n = 1-2; Fig. 2). LGDs were considered to have medium effectiveness (26–75%; n = 3; Fig. 2), while predator removal was deemed ineffective, as attacks were not reduced after culling of wolves, although this was evaluated only in a single case (Fig. 2).

#### Perceived effectiveness

For perceived effectiveness, two types of DMMs lacked data (improvement of agricultural practices and predator removal; Fig. 2). Information dissemination obtained the highest number of reports (n = 32), followed by electric fences, which were evaluated 27 times. Perceived effectiveness for the remaining DMMs was reported less frequently, with 10 or less projects reporting for each method.

DMMs perceived as being most effective included electric fences, LGDs, shepherds and visual/sound deterrents. Physical barriers, preventing access to anthropogenic food sources, and damage compensation were perceived as effective, while the rest were considered to be non-effective or giving mixed results (Fig. 2).

## Discussion

LIFE projects invest considerable resources in supporting wildlife management and conservation actions in Europe, including actions aimed to address human-wildlife conflicts. Our overview revealed considerable bias in the distribution and number of projects targeting large carnivores in certain parts of Europe, which does not reflect current distribution of large carnivores on this continent (Chapron et al., 2014; Cimatti et al., 2021). We observed a clear north-south gradient, with a very small number of projects in northern Europe, although these countries harbor important proportion of entire European populations of large carnivores. Most LIFE projects aimed to mitigate damages caused by these species have been conducted in the Mediterranean countries (Portugal, Spain, Italy and Greece) and Romania. We also observed a bias between eastern and western countries, with a lower number of projects conducted in the East (Romania being a notable exception). Reasons for observed bias in the spatial distribution of LIFE projects are not evident from data available, but we assume that the lower number of projects in northern countries might be connected with higher national budgets available for large carnivore management, research and conflict prevention in these countries, compared to Southern and Eastern Europe, where the need for EU funds is higher. A lower number of projects in Eastern Europe compared to Western Europe could be partly explained by their later accession to the European Union. A large number of LIFE projects in Romania might be connected with large populations of three species of large carnivores (wolf, bear, and Eurasian lynx) in this country (Chapron et al. 2014). An exceptionally high number of projects in Italy and Spain can be partly explained by several LIFE projects developed during the 1990’s, which were divided in two- or three-phased projects. Although they had the same goals and location, each phase was officially a separate project. Nevertheless, even if we had considered such multi-phase projects as a single project, Italy and Spain would still stand out as countries with the highest number of LIFE projects on large carnivores.

Our overview also revealed considerable differences between the numbers of projects dedicated to each species, especially a low number of projects dedicated to mitigation of damages caused by Eurasian lynx. Eurasian lynx is the least numerous among the treated taxa in Europe (Chapron et al. 2014), and is generally causing considerably less conflicts compared to wolf or bear (Bautista et al., 2019; Kaczensky, 1999), therefore there was probably less need for the application of DMM for this species, which is reflected in lower number of such projects. An exception is Northern Europe, where damage caused by lynx is frequent, especially where sheep graze without any protection (Odden et al., 2008). However, as noted above, number of LIFE projects was generally low in this region.

Despite the considerable funds provided through the LIFE program to EU Member States to help mitigate damages caused by large carnivores to human property, several limitations surfaced when we tried to measure their actual impact on economic losses to carnivores. This was mainly related to the availability and quality of the reported data. First, only 30 out of 74 projects made the final reports available, which limited the amount of information we could gather about the methods applied, their effort and effectiveness. The number of reports available was particularly low for the projects conducted in the 1990’s, as most of the documents were lost over the years. Therefore, the only information available from these projects was that provided in the project summary from the LIFE webpage, which was fairly limited, especially concerning the effort of each DMM and their effectiveness. Second, when the project reports were available, we observed that standards of measuring functional effectiveness were in general relatively low, with considerable inconsistencies in measurements. They were mostly limited to before-after comparison with non-random selection of treated locations, which does not provide strong inference when evaluating success of DMMs (Treves et al. 2019). This is mainly connected with several potential confounding factors that could affect the results and potential biases involved in evaluating effectiveness of interventions. For this reason, researchers recommend using at least a quasi-experimental approach, which in addition to treated locations or objects also includes randomly-selected controls (Treves et al., 2019). In most cases it was also not possible to discern potential biases involved in selection, treatment, measurement, and reporting. Furthermore, LIFE projects rarely reported the duration of the effect, although this is a crucial parameter that can considerably limit the success of DMMs, especially when long-term solutions are attempted to achieve (Khorozyan & Waltert, 2019). Although some DMMs, such as those involving pain stimuli (e.g. electric fences), can provide long-lasting solutions, effectiveness of others (e.g. various deterrents) often erode after few months (Khorozyan & Waltert, 2019).

Because of these limitations, our capacity to properly evaluate many of the used methods and provide sound guidelines was greatly limited. This limits the ability to adapt methods and increase success of conflict prevention in the most important conservation program in the European Union. However, this issue is not limited to LIFE projects, and recent reviews have pointed out a general lack of reliable information on the effectiveness of methods used to mitigate the damage caused by large carnivores (Treves et al., 2016, 2019; Eklund et al., 2017; Van Eeden et al., 2018).

In spite of the limitations presented above, we believe that the data on functional and perceived effectiveness of used DMMs does provide certain insights that could benefit future endeavors to reduce conflicts between people and large carnivores. The first conclusion is that most of the non-lethal methods showed at least moderate effectiveness (judged using both functional and perceived effectiveness), which is in agreement with previous studies (Berzi et al., 2021; Khorozyan & Waltert, 2020; Salvatori & Mertens, 2012; Treves et al., 2016; van Eeden et al., 2018). Sample size for several of these methods was, however, too small to be able to recommend them for widespread use in the future before more data becomes available. The only method that was tested more frequently (n=13) and was mostly reported as being highly effective, are electric fences. Based on this, it appears safe to recommend continued use of electric fences also for future projects, which also supports previous research done on the functional effectiveness of this method (Van Eeden et al. 2018; Khorozyan & Waltert, 2019, 2020). The sample size for actions related to the lethal removal of large carnivores evaluated within the LIFE program was too small and limited to only one species (grey wolf) and one case, as to be able to make any generalizations.

Several other factors beside the type of DMM used can affect occurrence of damage caused by carnivores on human property. One of the crucial ones is appropriate use and application of these measures (Frank & Eklund, 2017). For instance, electric fences require regular maintenance in the field to ensure their effectiveness. Some projects reported reduced effectiveness when fences were damaged or not set properly (e.g., improper grounding or too small fence circumference in respect to the sheep herd size or number of beehives to be protected). Livestock guarding dogs can be quite effective as well, but require proper training and can be expensive (registration, veterinary assistance). Additionally, several projects reported incompatibilities between the shepherd and the animal, so they were returned to the breeder. This points to the importance of close partnership with the livestock owners and communication on how to properly maintain DMMs, which was essential part of several successful LIFE projects.

Social and environmental context can also affect the effectiveness of measures used to prevent human-carnivore conflicts (Graha et al., 2005; Krofel et al. 2020). For example, availability of natural prey can be a critical factor, as reduced abundance might increase predators’ attempts to approach human settlements or livestock, although areas with higher prey density might also attract predators or increase their density, sometimes leading to increase in attacks on livestock (Janeiro-Otero et al., 2020; Treves et al., 2004). Other factors reported as important in previous research include the effects of surrounding habitat (e.g. distance to forest; Treves et al., 2011), meteorological conditions (Towns et al., 2009), breed of domestic animals (Bassi et al., 2021), herd size (Dar et al., 2009), status of local carnivore populations (e.g. stability of wolf packs; (Imbert et al., 2016; Santiago-Avila et al., 2018), historic presence of large carnivores in the area (Linnell, 2013), carnivores’ space use (Melzheimer et al., 2020) and their habituation to human presence (Majić & Krofel, 2015). Due to small sample sizes and limited availability of such information in LIFE project reports, we were not able to evaluate these effects on the effectiveness of prevention measures used in these projects. Nevertheless, these aspects should not be disregarded in future research, when more data becomes available.

### Recommendations for future projects

Considering the current limited amount of information and the high investments made across Europe under the LIFE program for reducing damage caused by large carnivores to human property, we call for higher standards in measuring effectiveness of DMM within LIFE projects, and similar programs. This will be especially important with the evaluation of novel, untested methods that might be applied in the future.

Specifically, our main recommendations for the future LIFE and other similar projects that include conservation actions aimed at mitigating damages to human property are: 1) provide clear methodology how functional and perceived effectiveness was evaluated; 2) prioritize measuring functional effectiveness, which is superior to measures of perceived effectiveness because of the problem of the placebo effect and the problem of researcher bias in encouraging positive responses by participants (Ohrens et al. 2019); 3) include randomly-selected control sites, where damages are measured in the same way as at treated sites, if possible with cross-over design (Treves et al., 2019); 4) report effectiveness for each DMM separately; 5) clearly indicate the investment made associated with every DMM (i.e. report budget, number of farms included, etc.); and 6) report any potential selection, treatment, measurement or reporting biases (Treves et al., 2019). We believe these recommendations would represent a major benefit for future LIFE projects, as well as other efforts of reducing conflicts between people and carnivores.

Additional information that would be useful to enable better evaluation of potential factors influencing effectiveness of implemented methods, include population status of large carnivores in the area (including pack stability in case of wolves), historic presence of large carnivores in project area (e.g. whether the area was recently recolonized or populations have never been exterminated), prey availability in project area, habitat characteristics (especially distance to forest for treated and control sites), and time period over which given method was applied and effects were measured.

We also advise more efforts focused on regions and species that have been so far largely neglected in LIFE projects, especially where funding from other sources is limited or when the complexity of bureaucratic procedures is deterring the owners from requesting assistance in damage prevention (Berzi et al., 2021). This includes several parts of the Central and Eastern Europe, and areas where damages caused by the Eurasian lynx represent significant issue locally.

## Supporting Information

Categories of DMMs and definition for each category is provided in Appendix S1.

## Acknowledgments

We are grateful to the teams of EuroLargeCarnivores Project and NEEMO Monitoring, especially R. Hickisch and J. Sliva, for their help in providing useful contacts and information on LIFE projects, which helped us to improve the database, especially for older projects. The study was funded by the European Union as part of the EuroLargeCarnivores LIFE project (grant number LIFE16 GIE/DE/000661). JVLB was additionally supported by the Spanish Ministry of Economy, Industry and Competitiveness (RYC-2015-18932; CGL2017-87528-R AEI/FEDER EU). MK was additionally supported by the Slovenian Research Agency (grants P4-0059 and N1-0163).

## Notes

### Competing Interest Statement

The authors have declared no competing interest.

