## Appendix S1 for "The contribution of the European LIFE program to mitigate damages caused by large carnivores in Europe"

**Supporting Information**

**Appendix S1.** Definitions for each DMM defined, largely following classification proposed by Treves et al., 2009.

| DMM | Definition |
| --- | --- |
| Large Carnivore Emergency Teams (LCET) | Teams of trained officers, with proper theoretical and practical training (capable of using a wide range of dissuasion tools including rubber-bullet rifles, tube traps, firecrackers, radio-collars and bear-dogs, as well as lethal removal of problem individuals), ready to act when a critical situation with a problem animal occurs (*e.g.* habituated bear in a village). |
| Information dissemination | Tools used to disseminate theoretical and technical information about the damage mitigation methods that can be used, or other activities that could potentially help reducing damages/or communication which we consider that will help to reduce damages. |
| Damage compensation schemes | A monetary compensation paid to a person which had part of his/her property damaged by large carnivores. |
| Visual and sound deterrents | Technologies used to disrupt large carnivores from unwanted behaviour/using given area. |
| Electric fences | Electrified wires or nets used to protect human properties (*e.g.* livestock or beehives). |
| Physical barriers | Physical barriers, such as stone, concrete or iron walls, to protect landowner’s property (*e.g.* livestock and beehives). |
| Preventing access to anthropogenic food sources | Methods used to prevent carnivores from accessing anthropogenic foods (e.g. garbage bins, trash, composts, slaughter remains) that can attract carnivores close to humans and cause further damage to their property. |
| Improvement of agricultural practices | Establishment of specific agricultural/farming practices compatible with large carnivore presence. (e.g. land rented to reduce livestock presence, improving monitoring and quality of sheep folds management, parasite control, improvement/creation of water sources). |
| Increasing food availability for carnivores | Increasing the amount of food available for wild large carnivores (i.e. natural or artificial supplementary food). |
| Livestock guarding dogs (LGDs) | Guarding dogs used to protect livestock from carnivore attacks. |
| Livestock guarding people | Employed staff or volunteers to help shepherds protecting livestock from large carnivore attacks. |
| Predator Removal | Culling or management removals of large carnivores to reduce damages. |
